# Cell-intrinsic Wnt4 controls early cDC1 commitment and suppresses development of pathogen-specific Type 2 immunity

**DOI:** 10.1101/469155

**Authors:** Li-Yin Hung, Yingbiao Ji, David A. Christian, Karl R. Herbine, Christopher F. Pastore, De’Broski R. Herbert

**Author notes:** These authors contributed equally (Karl R. Herbine, Christopher F. Pastore). Corresponding Author: De’Broski R. Herbert, Ph.D., School of Veterinary Medicine, University of Pennsylvania, Philadelphia, PA, 19104.

## Abstract

Whether conventional dendritic cell (cDC) precursors acquire lineage-specific identity under direction of regenerative secreted glycoproteins within bone marrow niches is entirely unknown. Herein, we demonstrate that Wnt4, a beta-catenin independent Wnt ligand, is both necessary and sufficient for necessary for the full extent of pre-cDC1 specification within bone marrow. Cell-intrinsic Wnt4 deficiency in CD11c^+^ cells reduced mature cDC1 numbers in BM, spleen, lung, and intestine and, reciprocally, rWnt4 treatment promoted pJNK activation and cDC1 expansion. Lack of cell-intrinsic Wnt4 in mice CD11c^Cre^*Wnt4*^flox/flox^ impaired stabilization of IRF8/cJun complexes in BM and increased the basal frequency of both cDC2 and ILC2 populations in the periphery. Accordingly, CD11c-restricted Wnt4 augmented Type 2 immunity against the hookworm parasite *Nippostrongylus brasiliensis* accompanied by increased interleukin 5 production relative to CD11c^Cre^ controls. Collectively, these data show previously unappreciated role for Wnt4 in DC commitment and pathogen-specific immunity.

## Introduction

Dendritic cells (DC) are professional antigen presenting cells that contribute to both innate and adaptive immunity through priming and instructing polarized T cell responses ^1^. Several types of DC can be found in both mouse and human, such as plasmacytoid DC (pDC), monocyte-derived DC (MoDC), Langerhans cells (LC) and conventional DC (cDC). Tissue-resident cDC are a diverse population that develops from a common myeloid progenitor (CMP) that gives rise to the macrophage and DC progenitor (MDP), which differentiates into a common DC progenitor (CDP) ^2^. FLT3-ligand drives conventional DC1 (cDC1) development from BM pre-cDC precursors under control of interferon regulatory factor 8 (IRF8) autoactivation that is stabilized through Batf3-Jun heterodimers forming a trimolecular complex required for cDC1 instead of cDC2 commitment ^3–7^. CD24 marks cDC1 that cross-present antigen to CD8^+^ T cells and serve as major producers of IL-12p70 for Th1 immune responses, whereas CD172a^+^ cDC2 are a heterogeneous population that require IRF-4 and promote Th2 and/or Th17 responses ^2,8^ Outside the transcriptional machinery required for DC-lineage commitment, it is less well known whether soluble proteins in BM niches control CDP lineage specification into cDC1 vs. cDC2 subsets.

Secreted glycoproteins in the mammalian Wingless/Integrase (Wnt) family serve important roles in embryonic development, cancer biology, and tissue regeneration through regulating cell proliferation and cell fate specification ^9–11^. Wnt signaling drives transcriptional activity through either canonical beta catenin-dependent transactivation or a non-canonical planar cell polarity (PCP) pathway that relies upon Jnk kinase activity ^12^. Wnt activation exerts immunoregulatory properties that remain poorly defined and somewhat contradictory ^13^. CD11c-driven deletion of β-catenin impairs Treg function and increased Th1/Th17 responses ^14,15^, but constitutive activation of β-catenin in CD11c^+^ cells also enhances Th1 responses ^16^. Exogenous addition of rWnt protein to DCs *in vitro* can change cytokine production profile ^14,17,18^ and within a tumor microenvironment, melanoma-derived paracrine Wnt5a promotes tolerogenic DC activity through metabolic reprogramming ^19^. Regarding non-canonical Wnts, *Wnt4* overexpression in hematopoietic stem cells (HSC) increased the frequency of myeloid progenitors and FLT3^+^ HSCs ^20^ and also increased numbers of fetal hematopoietic progenitors through Rac1 and JNK-dependent mechanisms ^21^, but overall, the role for Wnt production and signaling in DC biology is poorly understood.

In this study, CD11c^Cre^Wnt4^flox/flox^ mice were produced to test the cell-intrinsic role for Wnt4 in cDC biology during bone marrow differentiation, tissue homeostasis, and pathogen infection. Wnt4 deficiency disrupted the balance of pre-cDC 1 vs. pre-cDC2 in BM and preferentially reduced the numbers of mature cDC1 in the small intestine. This reduced cDC1/cDC2 ratio was accompanied by enhanced IL-33 receptor (ST2)-expressing ILC2, increased interleukin-(IL-) 5 production, and augmented immunity against the hookworm parasite *Nippostrongylus brasiliensis*. Taken together, these data support a model wherein Wnt4 acts in a cell-intrinsic manner in CDP to promote development of cDC1 in BM niche(s).

## Materials and Methods

### Mice

CD11c^Cre^(B6.Cg-Tg(Itgax-cre)1-1Reiz/J) and *Wnt4*^flox/flox^ (B6;129S-*Wnt4^tm1.1Bhr^*/BhrEiJ) mice were purchased from JAX laboratory and interbred to homozygosity. Age and gender matched CD11c^Cre^ and *Wnt4*^flox/flox^ strains were used as WT controls. Mice were housed under specific-pathogen free barriers in vivarium at San Francisco General Hospital or University of Pennsylvania School of Veterinary Medicine. All the procedures were reviewed and approved by IACUC at University of California at San Francisco (protocol #AN109782-01) and University of Pennsylvania (protocol #805911).

### Single-cell RNA Sequencing

Bone marrow cells were isolated and red blood cells were lysed before subjecting cells to CD11c MicroBeads (Miltenyi Biotec, Germany) for enrichment. Single-cell RNA sequencing was performed using 10X Chrome platform. The sorted cells were loaded onto a GemCode instrument (10x Genomics, Pleasanton, CA, USA) to generate single-cell barcoded droplets (GEMs) according to the manufacture’s protocol using the 10x Single Cell 3’ v1 chemistry. The resulting libraries were sequenced across two lanes on an Illumina HiSeq2500 instrument in High-output mode. Reads were aligned and subsequent analyses performed using the Cell Ranger (Pipeline). We obtained ~42k (36k form control and 51k form CD11c^Cre^-Wnt4^flox/flox^) reads per cell with a median genes per cell of 715 for control and 957 for CD11c^Cre^-Wnt4^flox/flox^ and median UMI count per cell of 2,639 for control and 2,832 for CD11c^Cre^-Wnt4^flox/flox^. The GEO accession number for the scRNA-seq data is

### Flow cytometry and PrimeFlow

Flow cytometry were performed on LSRII or LSRFortessa (BD Biosciences, San Jose, CA) with Diva software (version 8.0 or 8.0.1). Analysis was done in FlowJo 9.9.6 (FlowJo LLC, Ashland, OR). PrimeFlow^TM^ kit was purchased from Affymetrics (Santa Clara, CA) and the ViewRNA^®^ probes for mouse *Actb*, *Wnt4* and *Wnt16* were also designed and generated by Affymetrics. The assay was performed following the manufacturer’s protocol. The following flow antibodies were purchased from Biolegend (San Diego, CA): CD45 (30-F11), CD115 (AFS98), CD117 (2B8), CD135 (A2F10), CD11c (N418), MHCII (M5/114.15.2), Siglec-H (551), CD172a (P84), Ly6C (HK1.4), CD24 (M1/69), CD64 (X54-5/7.1), F4/80 (BM8), B220 (RA3-6B2), CD3 (145-2C11), Ter-119 (TER-119), Ly6G (1A8), CD19 (1D3), NK1.1 (PK136), XCR1 (ZET), CD103 (2E7), CD8a (53-6.7), CD11b (M1/70), Thy1.2 (30-H12), ICOS (15F9), CD127 (A7R34), ST2 (D1H9), GATA3 (16E10A23), IL12-p40 (C15.6). From Tonbo Biosciences (San Diego, CA): CD4 (GK1.5). From eBioscience (Santa Clara, CA): Gr-1 (RB6-8C5), CD5 (53-7.3), CD26 (H194-112). The live/dead fixable Aqua dead cell dye was purchased from Invitrogen (Carlsbad, CA).

### Cell isolation and treatments

For BM cells, bones from all four limbs were collected and crushed using a sterilized mortar and pestle. Cells were then passed through 70 μm strainers into collection tubes for FACS staining. For Wnt4 add-back experiments, single cell suspensions were passed through Histopaque-1083 gradient (Sigma, St. Louis, MO). 500,000 cells were then plated out in IMDM cell culture media (10% FBS and 1% Penicillin/Streptomycin) containing 100 ng/ml Flt3L (Peprotech, Rocky Hill, NJ) for 7 days. Treatment groups also received 100 ng/ml recombinant mouse Wnt4 or 100 ng/ml R-Spondin (R&D Systems, Minneapolis, MN) or both. Equal volume of PBS was added to control wells. Spleen DC isolation was performed as described by Stagg *et al.* ^22^. In brief, spleens were digested with RPMI containing 2% FCS, 1 mg/ml Collagenase D (Roche, Indianapolis, IN), plus 20 μg/ml DNase I (Sigma) for 1h before passing through 100 μm cell strainers to obtain single cell suspensions. Lungs were finely minced and incubated in digestion buffer (DMEM with 0.15 mg/ml Liberase TL+ 14 μg/ml DNase I+ 0.4 mg/ml Dispase) for 1h at 37C on a shaker. Cells were extruded through 18G needles 3 times before passing through 70 μm strainers to obtain single cell suspensions. To isolate cells from the small intestine, approximately 10 cm of jejunum was collected and opened longitudinally. Fecal matters were removed by rinsing with PBS. Tissues were then incubated in HBSS+ 5 mM EDTA+ 2.5 mM DTT for 3x10 min at 37C on a shaker. At the end of incubation, tissues were moved to a fresh petri dish and finely minced, then incubated in digestion buffer (DMEM+ 2% FBS+ 0.05 mg/ml Liberase TM+ 0.02 mg/ml DNase I) for 50 min. Digested tissues were extruded through 18G needle and pass through 100 μm strainers to obtain single cell suspension.

Mesenteric lymph nodes were collected and incubated in RPMI with 2% FCS, 1 mg/ml Collagenase D, and 20 μg/ml DNase I for 20 min then physically dissociated with rubber syringe heads against 70 μm cell strainers. Five hundred thousand cells were plated out for re-stimulation with plate-bound anti-CD3 (clone 2C11) and anti-CD28 (clone 37.51, both from Biolegend) for 72h. Wells containing cells with RPMI-10 media only were used as controls.

### Phospho-JNK ELISA

Bone marrow cells were isolated and passed through Histopaque-1083 gradient. Cells were rested in IMDM media without FBS for 2h before treating with R-Spondin (100 ng/ml, R&D Systems), Wnt4 (100 ng/ml, R&D Systems), R-Spondin+ Wnt4 (100 ng/ml each) or vehicle (PBS). At each time point, cells were collected and washed once with cold PBS and lysed at 10^7^/ml in lysis buffer [1 mM EDTA, 0.5% Triton X-100, 5 mM NaF, 6M urea, 2.5 mM Na_4_P_2_O_7_, 1x Halt^TM^ protease inhibitor cocktail (Thermo Fisher, Waltham, MA), and 1x phosphatase inhibitor cocktail 2 (Sigma)]. Phospho-JNK ELISA kit was purchased from R&D Systems and assays were carried out following the manufacturer’s protocol.

### Co-immunoprecipitation (Co-IP)

Bone marrow cells were isolated and enriched with CD11c MicroBeads (Milteyni Biotec). Cells were washed twice with cold PBS and 5-10 million cells were lysed with 100 μL lysis buffer [10 mM Tris-HCl, 50 mM HEPES, 150 mM NaCl, 1% Triton-X100, and 1mM EDTA. pH 7.4, 1X Halt protease inhibitor (Thermo Fisher), and 1x phosphatase inhibitor cocktail 2 (Sigma)]. The protein concentration was measured using BCA assay kit (Thermo Fisher). Around 0.2 mg total protein was incubated with agarose-conjugated anti-cJun (G-4, Santa Cruz Biotechnology, Dallas, TX) or equal amount of agarose-conjugated normal mouse IgG (Santa Cruz Biotechnology) overnight at 4C. The IP complex were washed twice with PBS and run on 4-12% gradient gel (Invitrogen). After transfer the membranes were immunoblotted with anti-IRF8 (clone ZI003, 1:500, Invitrogen) followed by incubation with rabbit anti-mouse IgG (1:1000). Images were taken on ChemiDoc XRS^+^ (Bio-Rad, Hercules, CA) and densitometry analyses were performed using ImageJ.

### Nippostrongylus brasiliensis (N.b.) infection

Parasites were maintained on fecal culture incubated at 25C. Larvae from 7 to 14-day old cultures were collected and washed 3x in PBS with 1% Penicillin/Streptomycin. To infect, 650-700 L_3_ stage larvae per mouse were injected subcutaneously at the base of tail. Fecal pellets were collected between d6-8 from each mouse to assess adult worm fecundity through parasite egg count per gram of feces. On d9, mice were euthanized and the proximal half of small intestines were removed, opened longitudinally, and incubated in PBS for 2h at 37C before collecting and counting adult worms.

### RNA isolation and real-time qPCR

Approximately 0.5 cm of duodenum was removed for RNA extraction using NucleoSpin RNA Plus kit (Macherey-Negel, Dueren, Germany). For CD11c MACS-enriched BM cells, 2 x 10^6^ cells were used for RNA extraction. Five hundred nanograms of total RNA were used to generate cDNA with Maxima H Minus reverse transcriptase (Thermo Fisher). Quantitative real-time PCR was performed on the CFX96 platform (Bio-Rad). Gene expression levels were normalized to (*Gapdh*). Primers used: *Gapdh* FW 5’-AGGTCGGTGTGAACGGATTTG-3’, RV 5’-TGTAGACCATGTAGTTGAGGT-3’; *Il5* FW 5’-CTCTGTTGACAAGCAATGAGA-3’, RV 5’-TCTTCAGTATGTCTAGCCCCT-3’; *Ccl2* FW 5” -TAAAAACCTGGATCGGAACAAA-3”, RV 5”-GCATT AGCTTC AGA TTTA CGG GT - 3’.

### T cell differentiation and cytokine ELISA

Bone marrow progenitors were cultured for 7 days in RPMI complete media (with 10% FCS and 1% Pen/Strep) supplemented with 20 ng/ml murine GM-CSF (PeproTech) to differentiate DC. To prepare T cells, splenocytes and lymph node cells from OT-II mice were pooled and enriched using naïve CD4 T cell isolation kit (Miltenyi Biotec). On the day of experiment, DC were pulsed with 50 ug/well of OVA for 8h, washed, then co-cultured with T cells at a 1:10 (DC: T) ratio for 3 days. At the end of the initial differentiation, cells were harvested and re-stimulated with plate-bound anti-CD3 for another 72h before collecting supernatants. The anti-mouse IL-5 sandwich ELISA kit was purchased from eBioscience and all assays were performed following manufacturers’ protocols.

### Statistics

Statistical analyses were performed using GraphPad Prism version 7.0 (GraphPad, La Jolla, CA).

## Results and Discussion

### Loss of Wnt4 expression in CD11c-positive cells leads to change in DC within BM

Evidence that Wnt4 over-expression can drive the up-regulation of Flt3 expression in BM precursors ^20^, prompted us to speculate that this particular non canonical Wnt may regulate DC development. Therefore, CD11c^Cre^Wnt4^flox/flox^ mice were generated to broadly interrogate the role of Wnt4 in DC biology. We first asked whether there was any change in the landscape of the naïve BM compartment due to CD11c-driven Wnt 4 deficiency. A single-cell RNA-sequencing followed by analysis with Seurat package based on ~4,000 CD11c^+^ enriched BM cells was taken as an initial approach (Fig. 1A) ^23^. Data shown in Fig. 1B reveal 17 lineage-specific cell clusters in BM of CD11c^Cre^ mice including cDC1 and cDC2 populations, but only 14 lineage-specific cell clusters in CD11c^Cre^Wnt4^flox/flox^ strain with an absence of the *Xcr1 Batf3*, *Irf8* expressing cDC1 population. To further investigate, the datasets were re-aligned and gene ontology analysis was completed with GSEA ^24^. Focusing on the gene expression pattern within the cDC2 population that was equivalent between strains, we noted that cDC2 within CD11c^Cre^Wnt4^flox/flox^ BM were markedly more pro-inflammatory than CD11c^Cre^ controls with enhanced signature for both IFN-α/-γ responses and inflammatory response pathways (Fig. 1C). Increased *Ccl2* expression in CD11c enriched BM cells from CD11c^Cre^Wnt4^flox/flox^ mice compared to controls confirmed the sc-RNA seq analysis (Fig. 1D).

**FIGURE 1.**
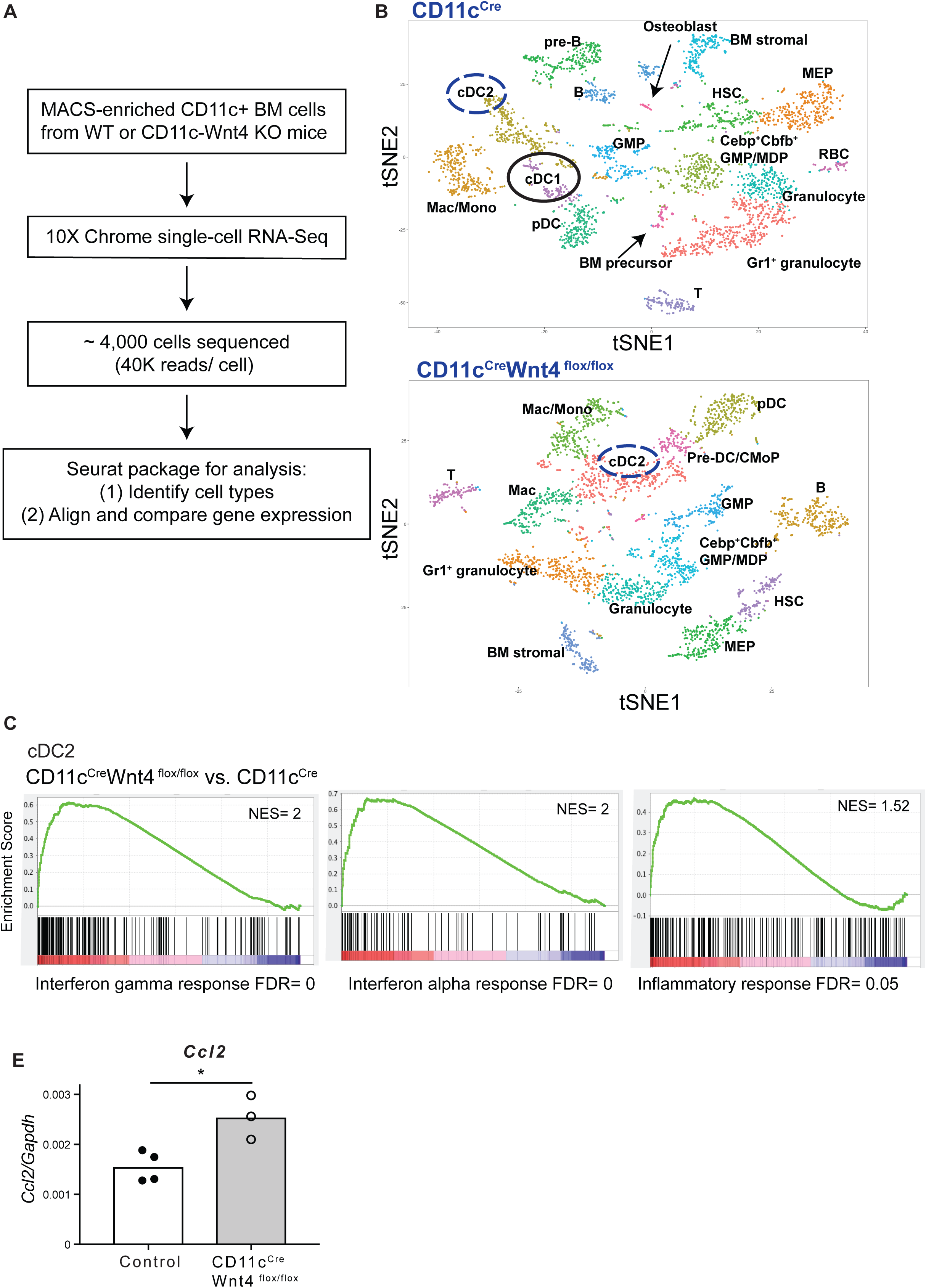
Single-cell RNA sequencing reveals altered cDC population and properties in CD11c^Cre^Wnt4^flox/flox^ BM. (A) Workflow of single-cell RNA-Seq and analysis. (B) Clusters in control (top) and CD11c^Cre^Wnt4^flox/flox^ (bottom) BM. (C) GSEA analysis of cDC2 population in BM. (D) Real-time PCR quantification of *Ccl2* levels in total CD11c^+^ BM cells from control and CD11c^Cre^Wnt4^flox/flox^ Representative results of 2 experiments. Graph shows Mean ± SEM with symbols represent individual mouse. *, p< 0.05 as determined by Students’ t-test.

### Wnt4 is expressed in multiple BM progenitor populations and controls the balance of pre-cDC1 vs. pre-cDC2

Given the lack of reliable mAb for Wnt4, we interrogated DC progenitors using PrimeFlow^TM^ assays to compare *Wnt4* mRNA expression levels and its downstream target *Wnt16* ^25^. Fig. 2A-B show that CD11c^Cre^ control and CD11c^Cre^-Wnt4^flox/flox^ BM expressed comparable *Wnt4* levels in the CDP population (CD135^+^CD115^+^CD117^med^CD11c^-^MHCII^-^) whereas *Wnt16* levels were un-detectable (Fig. 2A-C). Comparison of *Wnt4* levels in pre-cDC1 and pre-cDC2 subsets (SiglecH^-^CD135^+^CD117^med^CD172a^low^MHCII^low^CD11c^+^ and Ly6C^-^ or Ly6C^+^, respectively) ^26^ revealed significant reduction in pre-cDC1 within CD11c^Cre^-Wnt4^flox/flox^ compared to CD11c^Cre^ controls (Fig. 2C-2E). Intriguingly, the loss of *Wnt4* was most profound in a CD45^+^Lin^-^CD11c^+^CD115^-^MHCII^low^CD24^+^ population (Fig. 2F-G). CD24 is mainly expressed on pre-cDC1-primed, but not pre-cDC2-primed cells within DC progenitors ^26^, thus CD11c^Cre^-Wnt4^flox/flox^ BM had a lower percentage of pre-cDC1, but a relatively intact pre-cDC2 population (Fig. 2H-2I). Therefore, *Wnt4* was expressed in multiple DC progenitor cell types and its loss following CD11c-driven *Wnt4* depletion caused an imbalance of pre-cDC1/pre-cDC2 in the BM.

**FIGURE 2.**
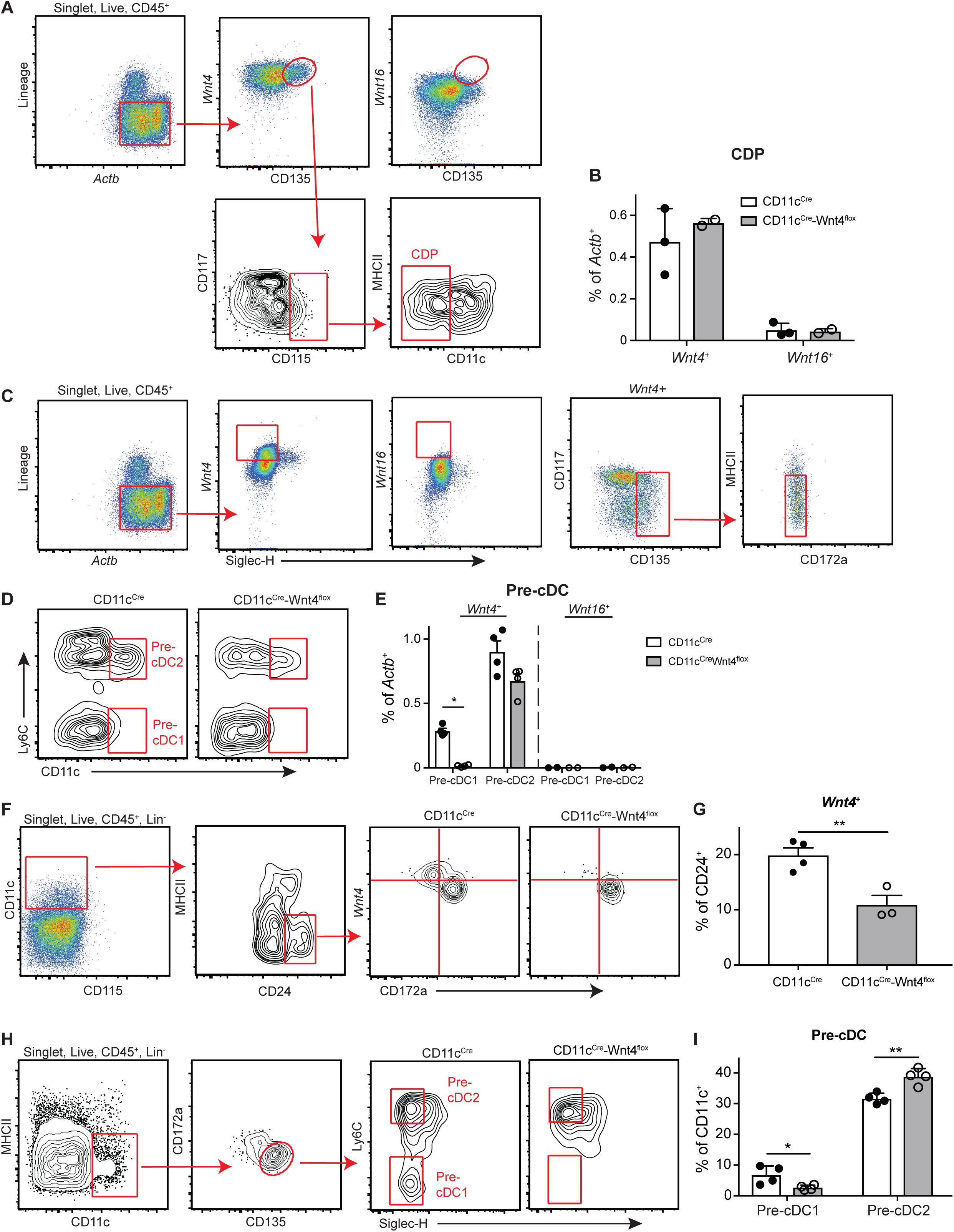
CD11c-driven Wnt4 deficiency affects cDC progenitors in BM. (**A**) Gating strategy to identify *Wnt4*+/*Wnt16*+ CDP population in BM. (**B**) Quantification of *Wnt4*^+^ and *Wnt16*^+^ CDP in CD11c^Cre^ vs CD11c^Cre^-Wnt4^flox/flox^ mice. (**C**) Gating strategy to identify *Wnt4^+^/Wnt16^+^* pre-cDC cells in BM. (**D**) Representative flow plots showing pre-cDC1 and pre-cDC2 populations in CD11c^Cre^ vs CD11c^Cre^-Wnt4^flox/flox^ BM after gated as in “**C**”. (**E**) Quantification of *Wnt4^+^/Wnt16^+^* pre-cDC1 and pre-cDC2 in CD11c^Cre^ vs CD11c^Cre^-Wnt4^flox/flox^ BM as gated in “**D**”. (**F** and **G**) (**F**) Gating and (**G**) percentage of *Wnt4*^+^ cell within the CD11c^+^CD115^-^CD24^+^ sub-population in BM. (**H** and **I**) (**H**) Gating strategy and representative flow plots from each strain and (**I**) quantification of pre-cDC1 and pre-cDC2 cells in BM from CD11c^Cre^ vs CD11c^Cre^-Wnt4^flox/flox^ mice. Lineage markers: B220, CD3, CD5, CD11b, CD19, NK1.1, Ly6G, and Ter-119. Representative results from 3 independent experiments. Graphs show Mean ± SEM with * p< 0.05 and ** p<0.01 as determined by Students’ t-test.

### Wnt4 promotes Jnk activation and IRF8 stabilization to promote cDC1 commitment

Even though CD11c-drestricted *Wnt4* deficiency disrupted pre-cDC1 commitment it remained unclear whether exposure of preDC to Wnt4 was sufficient to promote the cDC1 fate. To test this possibility, total BM from either CD11c^Cre^ or CD11c^Cre^Wnt4^flox/flox^ strains was cultured 7 days in the presence of Flt3L with either R-Spondin (R-Spd), rWnt4, or a combination of R-Spd and rWnt4. Upon harvest, both CD24^+^ (cDC1) and CD172a^+^ (cDC2) subsets were identified from the CD11c^+^MHCII^+^ population (Fig. 3A-B). Data show addition of rWnt4 or R-Spd^+^ alone moderately increased CD24^+^ percentage and cell number as compared to mock-treated CD11c^Cre^ controls, but that combined Wnt4/R-Spd treatment led to a 3-fold increase in CD24^+^ cells within 7 days. On the other hand, Wnt4/R-Spd treatment only partially expanded CD24^+^ cells in BM cultures of CD11c^Cre^Wnt4^flox/flox^ mice and had no effect on BM cultures from Batf3KO mice (Fig 3B-D). Regarding CD172a^+^ cells, although were no significant differences in percentage between CD11c^Cre^ and CD11c^Cre^Wnt4^flox/flox^ strains, the CD172a^+^ population was more evident in the latter following 7 days of combined Wnt4/R-Spd treatment, something that also increased the number of CD172a^+^ cells (Fig.3E-F). This suggests that while exogenous Wnt4 can promote cDC differentiation, either through driving cell proliferation or preventing cell death, the effect likely requires autocrine Wnt4 production.

**FIGURE 3.**
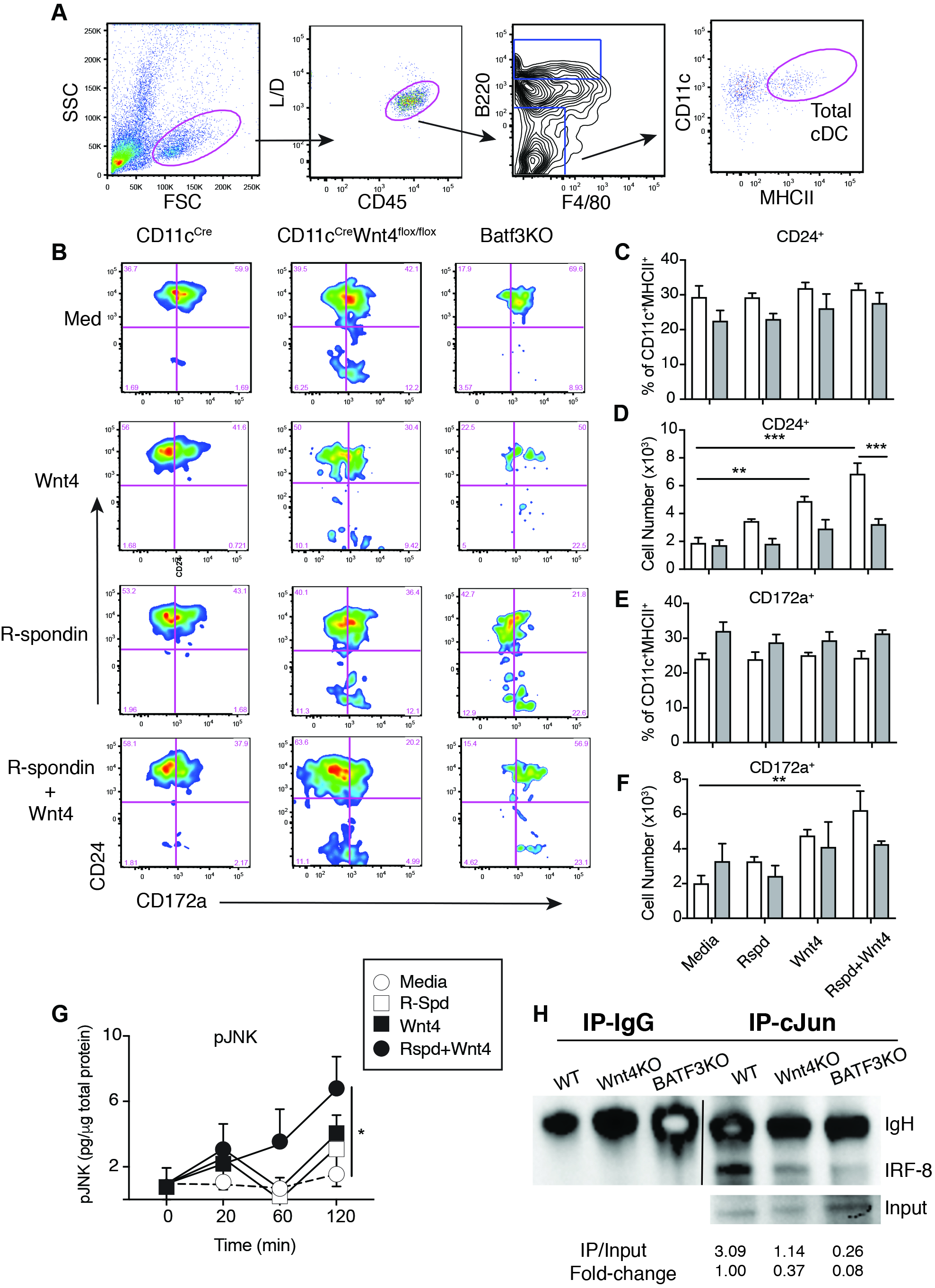
Wnt4 promotes differentiation of cDC, pJNK phosphorylation and cJun-IRF8 complex formation. (**A**) Gating strategy to identify cDC in Flt3L-induced BMDC cultures. (**B**) Representative flow plots after gated as in “**A**” from CD11c^Cre^ and CD11c^Cre^-Wnt4^flox/flox^ BM cells cultured for 6 days in IMDM CM+ 100 ng/ml Flt3L supplemented with R-Spd (100 ng/ml), Wnt4 (100 ng/ml), R-Spd+ Wnt4 (100 ng/ml each), or vehicle (PBS). (**C-F**) Quantification of (**C** and **E**) frequencies and (**D** and **F**) cell number of CD24^+^ (**C**, **D**) and CD172a+ (**E**, **F**) from conditions described in “**B**”. (**G**) Quantification of phospho-JNK at indicated time points from total BM cells after isolation and Histopaque enrichment, 2h serum starvation and treated with IMDM CM alone or media containing R-Spd (100 ng/ml), Wnt4 (100 ng/ml), or R-Spd+ Wnt4 (100 ng/ml each). (**H**) Representative gel picture showing co-immunoprecipitation of cJun and IRF8 from CD11c^Cre^ vs CD11c^Cre^-Wnt4^flox/flox^ vs Batf3^-/-^ BM CD11c^+^ cells (top). Quantification of co-IP products after normalization to inputs were shown at the bottom. Graphs show Mean ± SEM from N=3-4 biological replicates per condition with * p< 0.05, ** p< 0.01, and *** p< 0.005 as determined by two-way aNOVa with Tukey correction for multiple comparison. Representative results from three experiments.

To delve into how Wnt4 was acting as a pro-cDC1 signal, we asked whether Wnt4/R-Spd treatment induced pJNK activation in BM progenitors as described in Wnt 4-treated stromal cells that undergo non-canonical Wnt signaling through the PCP pathway ^20,21^. Phosphorylation of JNK leads to activation of cJun, and AP-1 to form a hetero-trimeric complex with Batf3 and IRF8 to promote cDC1 differentiation ^6,27^. Thus, we postulated that Wnt4 functioned to recruit cJun for stabilization of IRF8 to enforce cDC1 commitment from nascent progenitors. Indeed, kinetic analysis showed that Wnt4/R-Spd induced sustained pJNK activity in BM cells (Fig. 3G). Additionally, co-immunoprecipitation experiments comparing CD11c^Cre^, CD11c^Cre^Wnt4^flox/flox^, and *Batf3* KO BM demonstrated that cJun bound avidly to IRF8 in CD11c^Cre^ BM, but this signal was 3-fold reduced in CD11c^Cre^Wnt4^flox/flox^ and nearly absent in *Batf3KO* (Fig. 3H). These results also indicate that binding of cJun to IRF8 is Batf3-dependent, which was previously untested. Taken together, our data indicates that Wnt4 activates JNK, promotes cJun/Batf3/IRF8 complex formation in CD11c^+^ BM progenitors to promote cDC1 differentiation.

### CD11c-driven Wnt4 expression is critical for cDC homeostasis

We reasoned that a basal imbalance of pre-cDC populations in BM would predispose CD11c^Cre^Wnt4^flox/flox^ mice towards disproportionate cDC1 and cDC2 numbers in peripheral tissues. Wnt4 deficiency reduced cDC1 percentages in both spleen and small intestine with relatively increased splenic cDC2 (Fig. 4A-4C). CD103^+^CD11b^-^ cDC are considered to be the main source of IL-12 production, although in mucosal organs they seem to have different functions in that colonic CD103^+^ DC suppress inflammatory responses through production of IDO1 and IL-18 binding protein; whereas CD103^+^CD11b^+^ cells are tolerogenic and promote Treg differentiation via retinoic acid ^28,29^. Because intestinal cDCs can be defined by different flow cytometry gating strategies ^30–32^, we additionally probed for cDC1 frequency using CD11b and CD172a to mark small intestine cDC2 ^33^. This approach revealed a selective reduction in the CD103 population within the CD11c^Cre^Wnt4^flox/flox^ mice (Fig. 3E-F), further supporting our contention that cell-intrinsic *Wnt4* deficiency disrupts cDC subset identity under homeostatic conditions.

**FIGURE 4.**
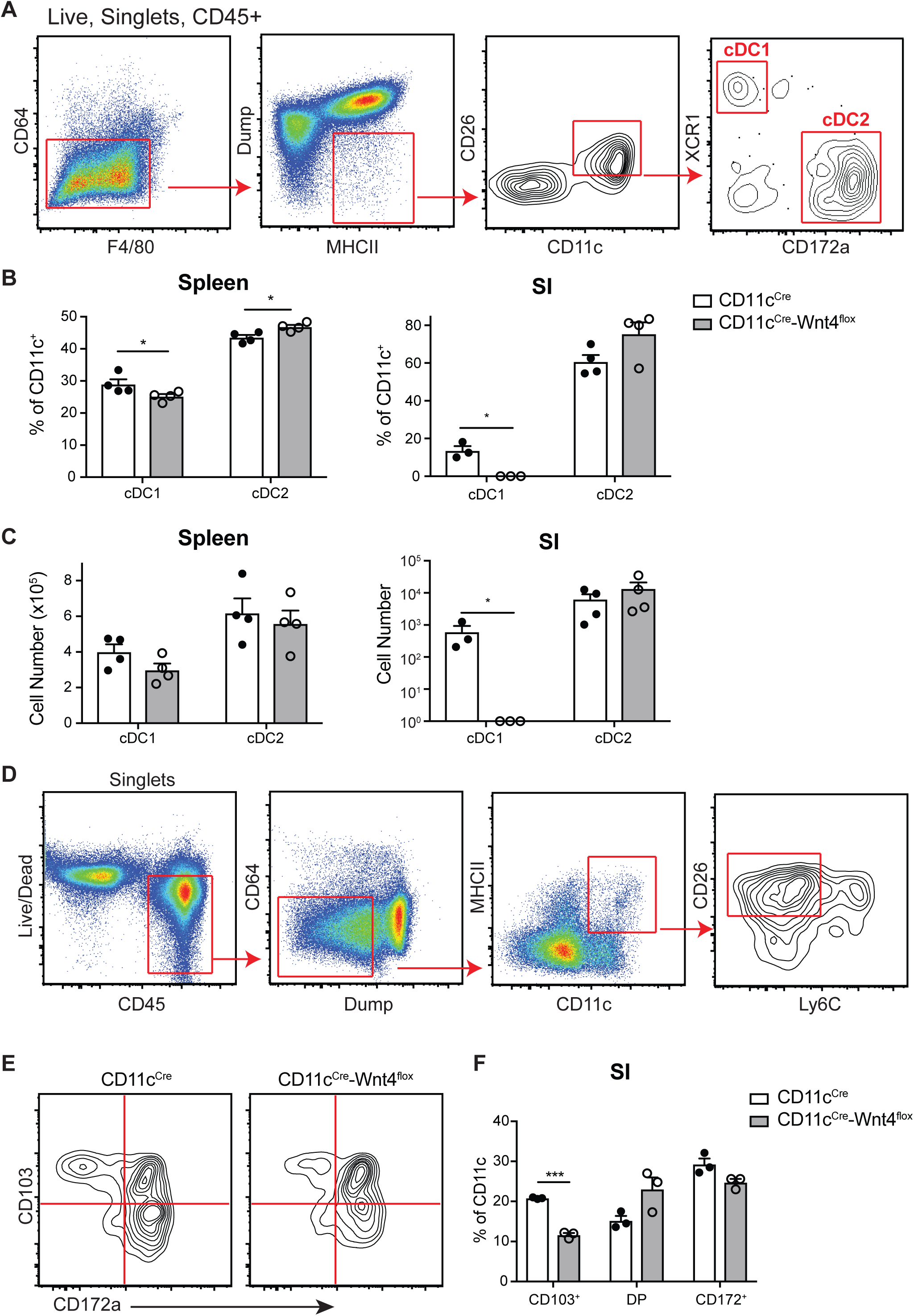
CD11c-specific Wnt4 deficiency affects cDC1 cell in periphery tissues at steady states. (**A**) Gating strategy to identify cDC1 and cDC2 in the spleen. (**B**) Percentage and (**C**) cell number of cDC1 and cDC2 in spleen and small intestine from CD11c^Cre^ vs CD11c^Cre^-Wnt4^flox//flox^ as identified in “**A**”. (**D**) Gating to identify cDC in small intestine. (**E** and **F**) (**E**) Representative plots and (**F**) quantification of cDC subsets in CD11c^Cre^ vs CD11c^Cre^-Wnt4^flox^ small intestine based on gating in “**D**”. Dump gate: B220, CD3, CD19, and NK1.1. Representative results from 3 to 4 independent experiments. Graphs show Mean ± SEM with symbols represent individual mouse. * p< 0.05 and *** p< 0.005 as determined by Students’ t-test.

### CD11c-mediated Wnt4 deficiency alters cDC distribution and causes lower helminth burden after Nippostrongylus brasiliensis (N.b.) infection

Because IL-12, a major cytokine produced by cDC1, inhibits host immunity against gastrointestinal nematodes, we tested whether Wnt 4 mediated dysregulation of intestinal cDC1 numbers would alter host protection against the hookworm *N. brasiliensis* (*N.b.*). In this model, inoculation of wild-type mice with 750 infectious stage larvae (L_3_), results in spontaneous clearance of adult parasites and termination of egg production between 9-12 days ^34,35^. Data in Fig. 5A shows CD11c^Cre^Wnt4^flox/flox^ mice had significantly lower fecal egg counts than CD11c^Cre^ controls between d6 through d8 post-infection. CD11c^Cre^Wnt4^flox/flox^ mice also eliminated their intestinal worms significantly faster than controls at both early (d4) and late (d9) stages of infection (Fig. 5B-C). This indicated that a basal dysregulation of the balance between cDC1 and cDC2 subsets altered the outcome of pathogen specific immunity.

**FIGURE 5.**
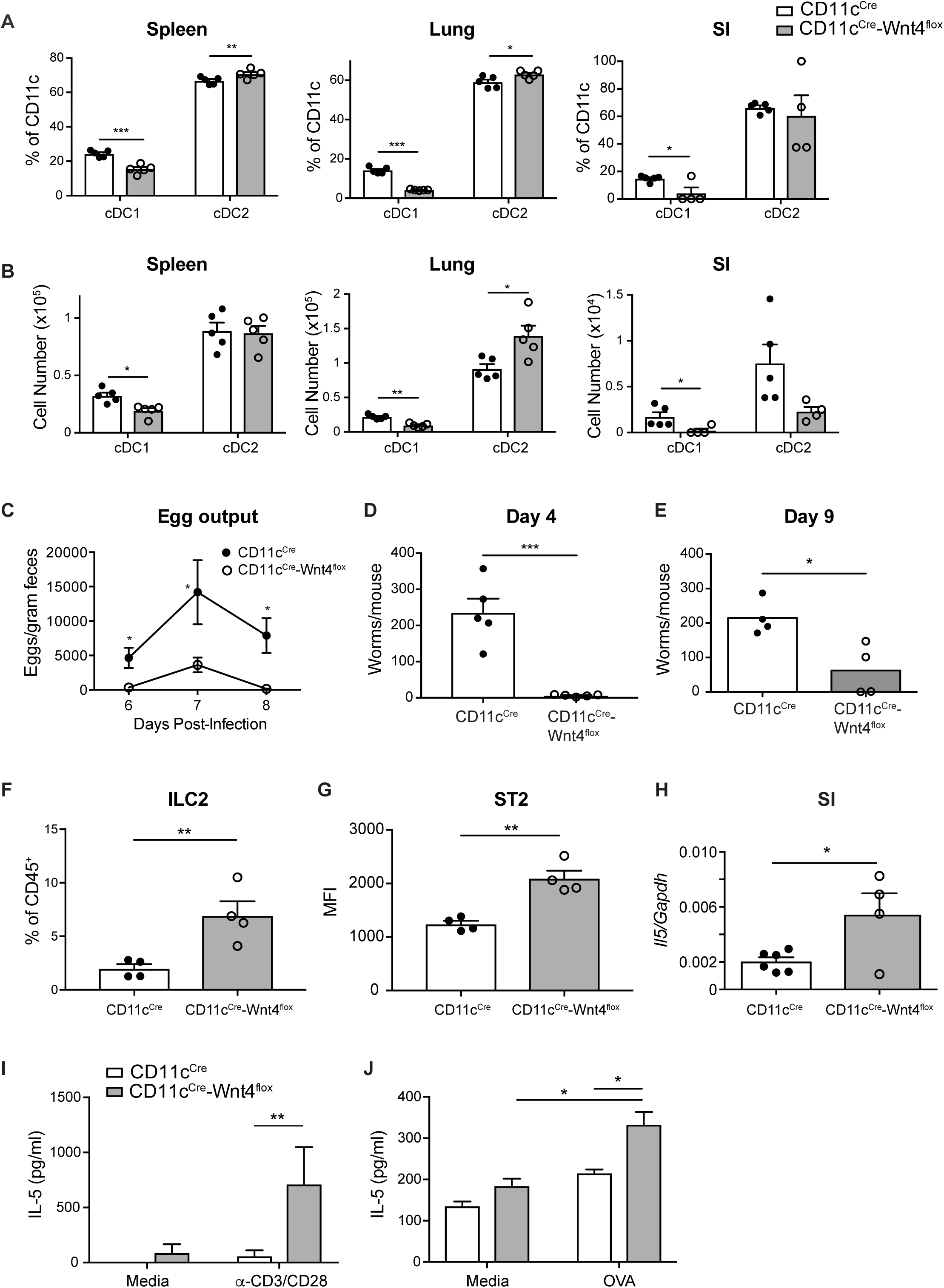
CD11c^Cre^-Wnt4^flox/flox^ animals mount stronger Th2 immune responses and accelerates worm clearance after *N.b*. infection. (**A**) Fecal egg counts on d6-8 from mice infected with 700 L_3_ *N.b*. larvae. (**B** and **C**) adult worm counts from CD11c^Cre^ and CD11c^Cre^-Wnt4^flox/flox^ mice on (**B**) d4 and (**C**) d9 after *N.b*. infection. (**D**) Percentage and (**E**) cell number of cDC1 and cDC2 in spleen, lung, and small intestines of CD11c^Cre^ vs CD11c^Cre^-Wnt4^flox/flox^ at d4 post-N.b. infection (700 L_3_ larvae). (**F**) Percentage of lung ILC2 and (**G**) Median fluorescence intensity (MFI) of ST2 expression on lung ILC2 from CD11c^Cre^ vs. CD11c^Cre^-Wnt4^flox/flox^ mice at d4 of *N.b*. infection. (**H**) Transcript levels of *Il5* mRNa in small intestine from CD11c^Cre^ vs. CD11c^Cre^-Wnt4^flox^ mince at d9 post-*N.b.* infection. (*I*) IL-5 levels from d9 *N.b.*-infected mesenteric lymph node cells after restimulating with media or plate-bound anti-CD3/CD28 for 72h (N=3-5/group). (**J**) IL-5 levels of OT-II T cells stimulated with OVa-pulsed CD11c^Cre^ or CD11c^Cre^-Wnt4^flox^ DC. DC cultured with media only were used as controls (N=3/group). Representative results from 2 to 3 independent experiments. Graphs show Mean ± SEM with * p< 0.05, ** p< 0.01, and *** p< 0.005 as determined by Students’ t-test or 2-way aNOVa with Tukey correction for multiple comparison.

To determine whether hookworm infection could reverse the basal defects in naïve CD11c^Cre^Wnt4^flox/flox^ mice through infection induced hematopoiesis we probed the lung, spleen and small intestines at day 4 post-*N.b.* infection. cDC1 were significantly reduced in both percentage and number within CD11c^Cre^Wnt4^flox/flox^ mice as compared to CD11c^Cre^ controls (Fig. 5D-E). This was accompanied by an increased percentage and number of cDC2 in lung and spleen, whereas the small intestine cDC2 population was similar between strains (Fig. 5D-E). Because a relative increase in cDC2 could enhance Type 2 responses, we evaluated both innate and adaptive sources of IL-5, a major product of group 2 innate lymphoid cells (ILC2) and Th2 cells, respectively ^36^. Both ILC2 percentages and ILC2 ST2 expression levels were significantly increased in CD11c^Cre^Wnt4^flox/flox^ compared to controls (Fig. 5F-G). Higher levels of ST2, the IL-33 receptor, implied increased sensitivity to the pro-type 2 alarmin cytokine IL-33 ^37^. Moreover, intestinal *Il5* mRNA transcripts and T cell mitogen (α-CD3)-stimulated IL-5 release from mesenteric lymph nodes of infected mice were both higher in CD11c^Cre^Wnt4^flox/flox^ compared to controls (Fig. 5H-I).

Lastly, to test whether DC lacking Wnt4 intrinsically promoted naïve T cells to become Th2 cells, we used a DC-T cell co-culture system that employed BMDC pulsed with ovalbumin and co-cultured with naïve OTII cells. Data show that OVA-pulsed CD11c^Cre^Wnt4^flox/flox^DC promoted greater levels of IL-5 as compared to CD11c^Cre^ BMDC counterparts (Fig. 5J). Collectively, this indicates that lack of Wnt4 in DC accelerates innate and adaptive Type 2 responses and anti-helminth immunity.

These data reveal that Wnt4 is an intrinsic regulator of DC development and homeostasis in mice. During kidney development, Wnt4 can be induced by morphogens such as bone morphogenic protein (BMP) and leukemia inhibitory factor (LIF) or environmental cues like high glucose ^38^. However, the cellular source and regulation of Wnt4 during hematopoiesis is still unclear. Wnt4 can activate either canonical or non-canonical Wnt signaling pathways depending on the context ^39–41^, but in HSCs, Wnt4 acts in a β-catenin-independent manner ^20^ and requires Frizzled 6 (Fzd6) to activate JNK2 and to induce the expression of c*Jun* and c*Fos* ^21^. Intriguingly, Fzd6 is highly expressed on in BM cells that are pre-committed to become cDC1 ^26^. However, whether Wnt4 signals through Fzd6 in either pre-cDC in BM or mature cDC or whether Wnt4 signaling regulates or interfaces with BATF3, a key transcriptional factor in cDC1 development ^42^ remain open questions. BATF3 forms heterodimers with JUN to partner with IRF8 to stabilize the early auto-activation of IRF8 during cDC1 development ^6,27^. In the absence of BATF3, the initial activation of IRF8 is not sustained and pre-cDC1 can reverse its developmental path to become pre-cDC2 ^6^. It is therefore possible that Wnt4 induction of *cJun* could affect pre-cDC commitment and subsequent cDC development through stabilizing the IRF8/BATF3 interaction.

We show that the deficit in cDC1 corresponded to augmented Type 2 responses and clearance of worm infection, which is consistent with reports showing that loss of IRF8-dependent cDC1 led to a strong Th2 response following low dose infection with the GI nematode *Trichuris muris* (*T.m.*) infection, whereas the lack of IRF4-dependent cDC2 delayed the clearance of *T.m.* ^43^. Evert *et al* demonstrated that the BATF3-dependent cDC1 subset is a critical source of IL-12, both at the steady state and after helminth infection ^44^. IL-12 is known to antagonize Th2 immune responses ^45^ and exogenous addition of IL-12 delays immunity against *N. brasiliensis* infection ^34^. Similar to CD11c-restricted Wnt4 deficiency, *Batf3* KO mice also cleared helminth infection faster than WT ^44^.

In summary, this report reveals a novel role for exogenous and cell-intrinsic Wnt4 as a critical regulator of cDC development in BM and peripheral organs. Loss of Wnt4 reduces cDC1 development and augments host immunity against hookworm infection. Future work is needed to interrogate whether Wnt4 effects on DC shapes the anti-tumor responses and cross presentation to CD8^+^T cells for immunity against tumors, viruses, or microbial pathogens.

## Acknowledgements

This work is supported by grants: UO1AI125940, AI095289, GM083204, and Burroughs Wellcome Fund.

## Author contributions

L.-Y.H, D.A.C, and D.R.H. conceived and designed experiments. L.-Y.H, D.A.C., Y.J., K.R.H., and C.F.P. performed experiments. L.-Y.H, Y.J. and D.R.H. analyzed data. L.-Y.H. and D.R.H. wrote the manuscript.

All authors reviewed the final manuscript. We thank Dr. Malay Haldar for critical comments. The authors declare no financial conflicts of interest.

